# Selective interferon responses of intestinal epithelial cells minimize TNFα cytotoxicity

**DOI:** 10.1101/2020.03.20.000604

**Authors:** Jacob A. Van Winkle, David A. Constant, Lena Li, Timothy J. Nice

**Affiliations:** Department of Molecular Microbiology and Immunology, Oregon Health & Science University, Portland, Oregon, USA

## Abstract

Interferon (IFN) family cytokines stimulate genes (ISGs) that are integral to antiviral host defense. Type I IFNs act systemically whereas type III IFNs act preferentially at epithelial barriers. Among barrier cells, intestinal epithelial cells (IECs) are particularly dependent on type III IFN for control and clearance of virus infection, but the physiological basis of this selective IFN response is not well understood. Here, we confirm that type III IFN treatment elicits robust and uniform ISG expression in neonatal mouse IECs and inhibits replication of IEC-tropic rotavirus. In contrast, type I IFN elicits a marginal ISG response in neonatal mouse IECs and does not inhibit rotavirus replication. *In vitro* treatment of IEC organoids with type III IFN results in ISG expression that mirrors the *in vivo* type III IFN response. However, the response of IEC organoids to type I IFN is strikingly increased relative to type III IFN in magnitude and scope. The expanded type I IFN-specific response includes pro-apoptotic genes and potentiates toxicity triggered by tumor necrosis factor alpha (TNFα). The ISGs stimulated in common by types I and III IFN have strong interferon-stimulated response element (ISRE) promoter motifs, whereas the expanded set of type I IFN-specific ISGs, including pro-apoptotic genes, have weak ISRE motifs. Thus, preferential responsiveness of IECs to type III IFN *in vivo* enables selective ISG expression during infection that confers antiviral protection but minimizes disruption of intestinal homeostasis.

## INTRODUCTION

Interferon (IFN) family cytokines provide antiviral defense through stimulation of a broad transcriptional response that includes direct-acting antiviral genes (1, 2). Members of the IFN family are divided into three types based on receptor usage: multiple type I IFN genes (many IFN-αs, IFN-β, others; hereafter IFN-α/β), a single type II IFN gene (IFN-γ), and multiple type III IFN genes (up to four IFN-λs) (3, 4). IFN-α/β and IFN-λ are produced upon detection of viral nucleic acids and are primary components of the early response to infection. The heterodimeric receptor for IFN-α/β (IFNAR) is expressed by most cell types, but the distinct heterodimeric receptor for IFN-λ (IFNLR) is preferentially expressed by neutrophils and epithelial cells (4, 5). Prior studies from us and others in mouse models of gastrointestinal virus infection have used receptor-deficient animals to show that IFN-λ is particularly important for protection of intestinal epithelial cells (IECs) (6–9). Additional mouse studies suggest that IECs require IFN-λ for antiviral protection because they are less responsive to IFN-α/β in comparison to other epithelial cell types (6, 10, 11), which may result from downregulated IFNAR expression *in vivo* (6, 11). However, the physiological benefit of this preferential IEC responsiveness to IFN-λ has remained unclear.

Activation of IFNAR or IFNLR results in phosphorylation of signal transducer and activator of transcription (STAT) transcription factors and upregulation of IFN-stimulated genes (ISGs). More specifically, STAT1 and STAT2 are phosphorylated, bind interferon response factor 9 (IRF9), and form a hetero-trimeric complex called interferon stimulated gene factor 3 (ISGF-3). ISGF-3 translocates to the nucleus and binds interferon-stimulated response element (ISRE) motifs in ISG promoters (1, 12, 13). Additionally, STAT1 homodimers, other STAT family members, and non-canonical factors also play a role in the transcription of some ISGs (14, 15). Prior comparisons of ISG induction by IFN-λ or IFN-α/β in cultured hepatocytes revealed largely overlapping responses consisting of canonical antiviral ISGs (16–21). However, other in-depth studies of neutrophils and hepatocytes have indicated that IFN-α/β is generally more potent than IFN-λ and results in greater chemokine and cytokine production (22–26). Additionally, studies of IECs cultured *in vitro* as 3D organoids have found that they are highly responsive to IFN-α/β, unlike IECs *in vivo* (27–32). Thus, the physiological basis of preferential IEC responsiveness to IFN-λ *in vivo* remains unclear.

Herein, we directly and quantitatively compare the IEC response to IFN-β and IFN-λ *in vivo* and *in vitro*. We find that the *in vivo* IEC response to IFN-β is minimal and does not inhibit replication of IEC-tropic rotavirus. In contrast, *in vitro* IFN-β treatment of IEC organoids elicits hundreds of ISGs, including pro-apoptotic genes, and potently blocks rotavirus infection. *In vitro* and *in vivo* IECs are equally responsive to IFN-λ and upregulate known antiviral genes but not pro-apoptotic genes. Consistent with differing pro-apoptotic gene expression, we show that cytotoxicity triggered by tumor necrosis factor alpha (TNFα) is increased in IEC organoids pre-treated with IFN-β relative to IFN-λ. Finally, bioinformatic scoring of promoter motifs indicates that IFN-β-specific ISGs, including pro-apoptotic genes, have low-scoring, weak ISREs. Antiviral ISGs stimulated in common by IFN-λ and IFN-β have high-scoring, strong ISREs. Together, these findings suggest that preferential responsiveness of IECs to IFN-λ *in vivo* ensures that antiviral ISGs are minimally accompanied by pro-apoptotic genes to promote epithelial homeostasis during clearance of enteric infection.

## RESULTS

### IECs in the neonatal intestine are minimally responsive to IFN-β

To extend our understanding of the transcriptional response to IFN in the intestine, we performed ISG *in situ* hybridization on intestinal tissues following IFN treatment *in vivo*. We injected PBS, IFN-β, or IFN-λ3 into seven-day-old neonatal mice, which have low baseline ISG expression, and detected transcripts for a canonical ISG (*Usp18*) four hours later. IFN-λ stimulated a robust increase in expression within the epithelial layer with no visible stimulation of cells in the underlying lamina propria tissue (**Fig. 1A**). In contrast, IFN-β injection resulted in a modest increase of *Usp18* in dispersed cells of the epithelium and lamina propria (**Fig. 1A**).

**Figure 1.**
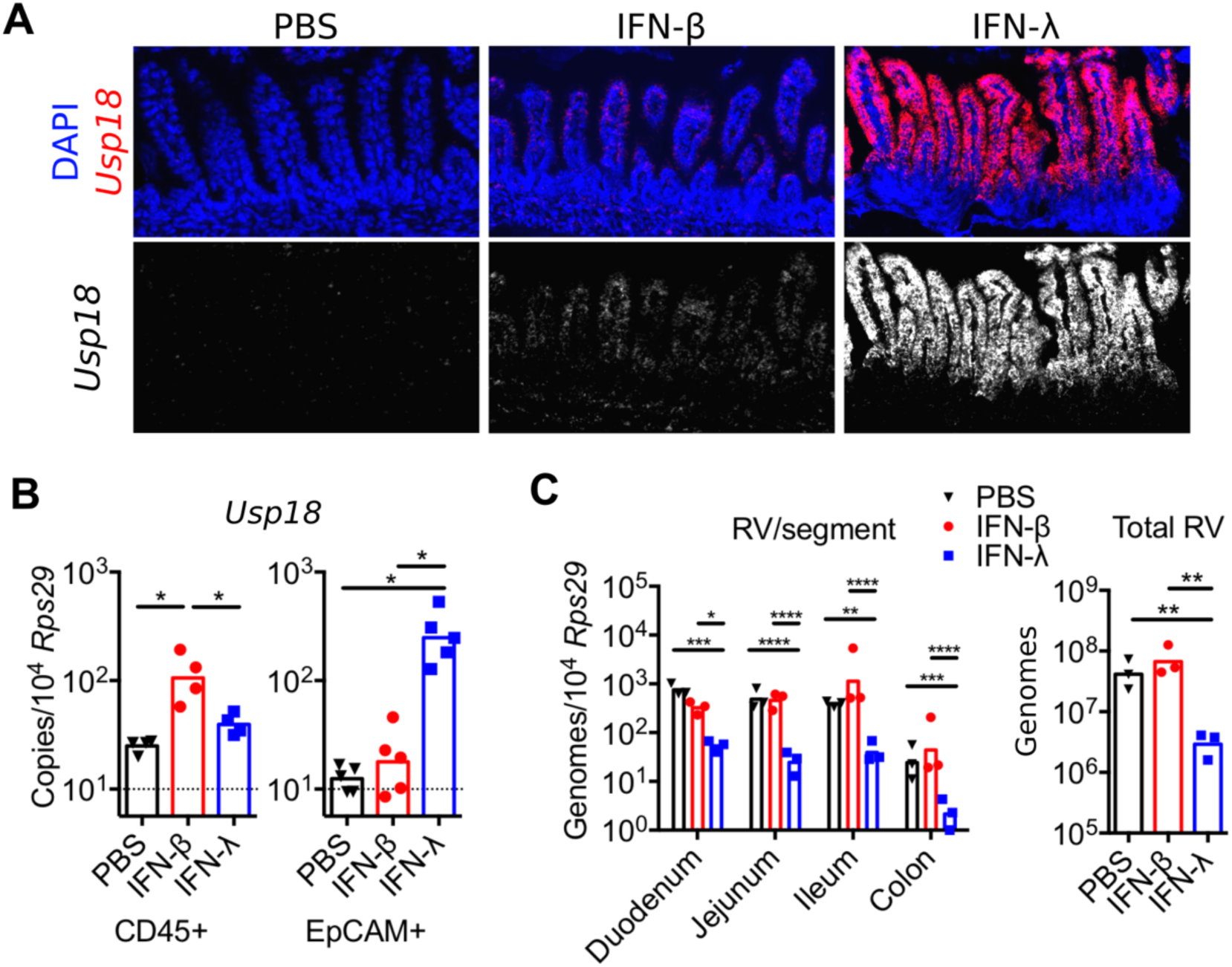
IECs in the neonatal intestine are minimally responsive to IFN-β. Neonatal mice were injected with either IFN-β or IFN-λ3 for four hours. **A.** Small intestinal tissue was isolated and stained with DAPI and *Usp18* antisense probes. **B.** *Usp18* abundance in sorted IECs (EpCAM-positive/CD45-negative) and hematopoietic cells (CD45-positive/EpCAM-negative) was analyzed by qPCR. **C.** mice were inoculated with rotavirus and viral genomes were quantitated in intestines 20 hours later. Data are combined from at least two experiments; individual replicates are shown with bars at mean values. Significance determined by one-way or two-way ANOVA; *, p < 0.05; **, p < 0.01; ***, p < 0.001, ****, p < 0.0001.

To more quantitatively compare transcript abundance in IECs and intra-epithelial hematopoietic cells, we FACS sorted EpCAM-positive/CD45-negative epithelial (EpCAM+) and CD45-positive/EpCAM-negative hematopoietic (CD45+) cells from dissociated epithelium. Consistent with *in situ* hybridization results, qPCR analysis showed that *Usp18* was stimulated more than 20-fold in EpCAM+ cells following IFN-λ injection but less than two-fold following IFN-β injection (**Fig. 1B**). Conversely, *Usp18* was stimulated less than two-fold in intra-epithelial CD45+ cells following IFN-λ injection but stimulated five-fold following IFN-β injection **(Fig. 1B**).

To determine whether *Usp18* transcripts were indicative of a broader ISG program that conferred antiviral protection upon IECs, we challenged neonatal mice treated as above with IEC-tropic murine rotavirus and quantitated viral genomes in the intestine 20 hours later. IFN-λ injection resulted in three- to ten-fold lower viral genomes compared to PBS injection, but IFN-β injection provided no significant protection (**Fig. 1C**). These data align with previous reports of the preferential IEC response to IFN-λ in adult mice, and suggest that hypo-responsiveness of IECs to IFN-α/β *in vivo* arises in early neonatal life.

### IEC organoids are dually responsive to IFN-β and IFN-λ

To determine whether IFN-α/β hypo-responsiveness was intrinsic to IECs, we generated *in vitro* IEC organoids from isolated epithelial stem cells (**Fig. 2A**). We stimulated these IEC organoids with 0, 0.1, 1, 10, or 100 ng/mL of mouse IFN-β or IFN-λ2 for 2, 4, 8, or 16 hours and quantitated the abundance of three canonical ISGs (*Isg15, Usp18*, and *Cxcl10*). All three ISGs were increased by IFN-β and IFN-λ2 treatments with a maximal upregulation between 100 and 1000-fold (**Fig. 2B**). The expression kinetics were similar for all three ISGs following IFN-λ stimulation, with maximal upregulation four to eight hours post-treatment and sustained expression at 16 hours. Similar expression kinetics were observed for *Isg15* and *Usp18* following IFN-β stimulation. However, at the highest doses of IFN-β, *Cxcl10* reached a peak of induction at four hours and decreased thereafter (**Fig. 2B**). Comparison of the dose-response for IFN-β or IFN-λ at four hours post-treatment indicated that early ISG upregulation was between two- and ten-fold greater for IFN-β compared to IFN-λ (**Fig. 2C**). These data indicate that IEC organoids upregulate canonical ISGs in response to IFN-β and IFN-λ, with minimal differences in expression kinetics, but higher maximum response to IFN-β. Therefore, IFN-α/β hypo-responsiveness observed *in vivo* is not an intrinsic property of IECs

**Figure 2.**
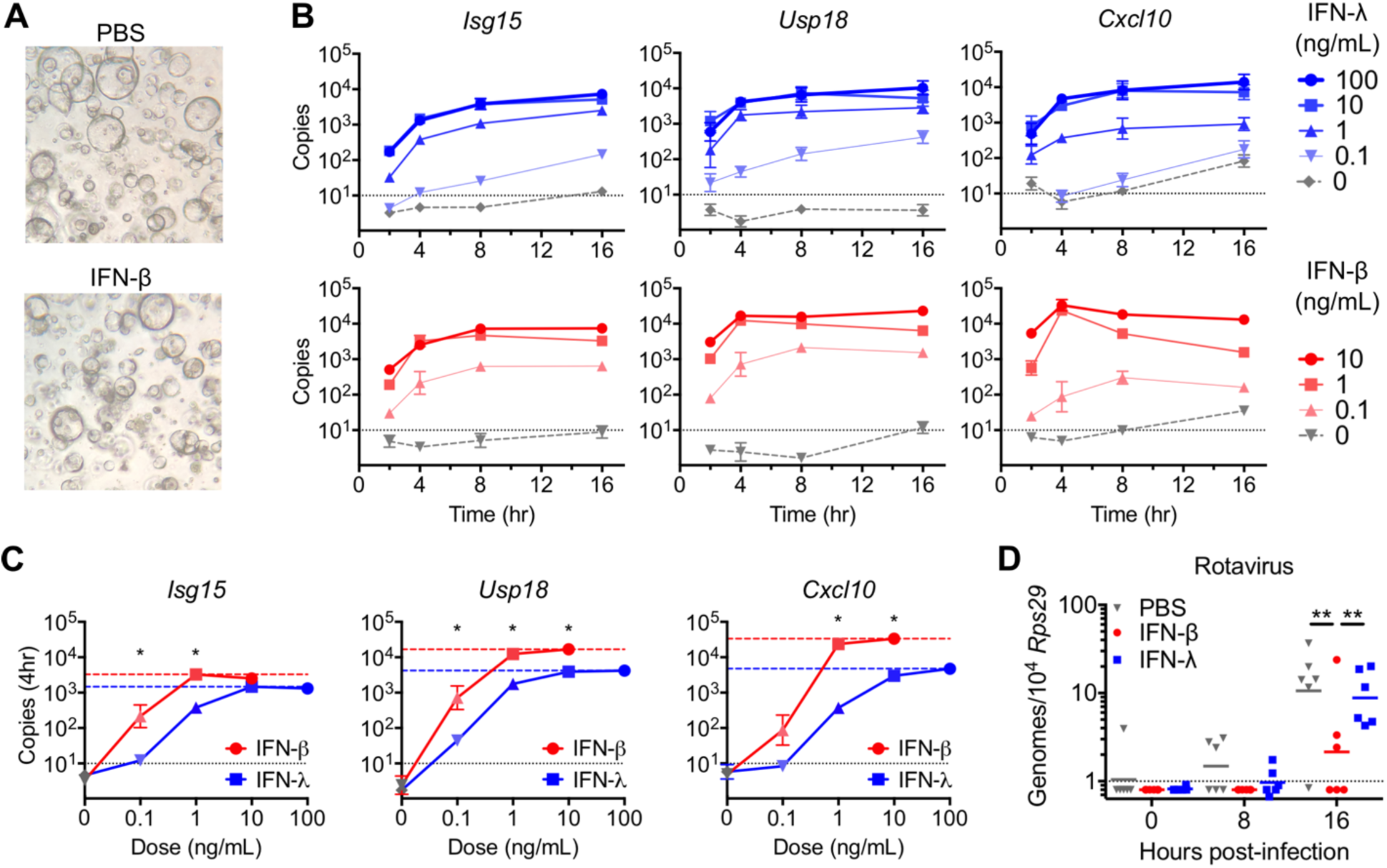
IEC organoids are dually responsive to IFN-β and IFN-λ. **A.** Representative images of IEC organoids. **B.** IEC organoids were treated with the indicated concentrations of recombinant IFN-β or IFN-λ2 for the indicated times and RNA was isolated for quantitation of *Isg15, Usp18*, and *Cxcl10* transcripts by qPCR. The dotted line indicates the limit of detection. **C.** Transcript abundance of the indicated genes four-hours after IFN treatment across the range of IFN doses tested. Dashed lines indicate maximal responses for IFN-β (red) or IFN-λ (blue). **D.** IEC organoids pre-treated with PBS, IFN-β or IFN-λ3 for eight hours were infected with rotavirus. Viral genomes were quantified at the indicated time post-infection. Data is from two (**B-C)** or three (**D**) experiments with duplicate treatment/infection wells; error bars show SEM. Statistical significance determined by t-test (**C**) or one-way ANOVA (**D**); *, p < 0.05; **, p < 0.01.

To confirm that the above ISG expression was indicative of the overall antiviral program stimulated by IFN treatments, we challenged IEC organoids with murine rotavirus and quantitated viral genomes 0, 8, and 16 hours later. IFN-λ treatment resulted in modest reductions in viral genomes relative to PBS treatment, but IFN-β treatment resulted in three- to ten-fold reduction in viral genomes (**Fig. 2D**). Therefore, IFN-β stimulates a stronger antiviral response than IFN-λ in cultured IEC organoids. These data indicate that the antiviral ISG response of IECs *in vivo* is regulated by factors not recapitulated in organoid culture.

### Global ISG expression in response to IFN-β is suppressed *in vivo*

To more comprehensively compare the *in vivo* and *in vitro* IEC response to IFN, we performed RNA sequencing on sorted EpCAM+ cells from neonatal mice treated with PBS, IFN-β, or IFN-λ3 and compared to RNAseq of IEC organoids treated with PBS, IFN-β, or IFN-λ2 for four hours. Normalized read counts for *Usp18* were increased between 5- and 1000-fold by IFN treatment, with a greater increase in IEC organoids by IFN-β treatment relative to IFN-λ treatment and a greater increase in neonatal IECs by IFN-λ treatment relative to IFN-β treatment (**Fig. 3A**). These differences are reflective of qPCR results from **figures 1-2**, validating the RNAseq dataset. PCA analysis indicated that the primary component differentiating these samples (PC1, 95% of variance) was their organoid or neonate origin and included differential expression of metabolism and cell cycle genes (**Fig. 3B, Data Set S1**). The secondary PCA component (PC2, 2% variance) separated IFN treatment groups from matched PBS controls (**Fig. 3B**).

**Figure 3.**
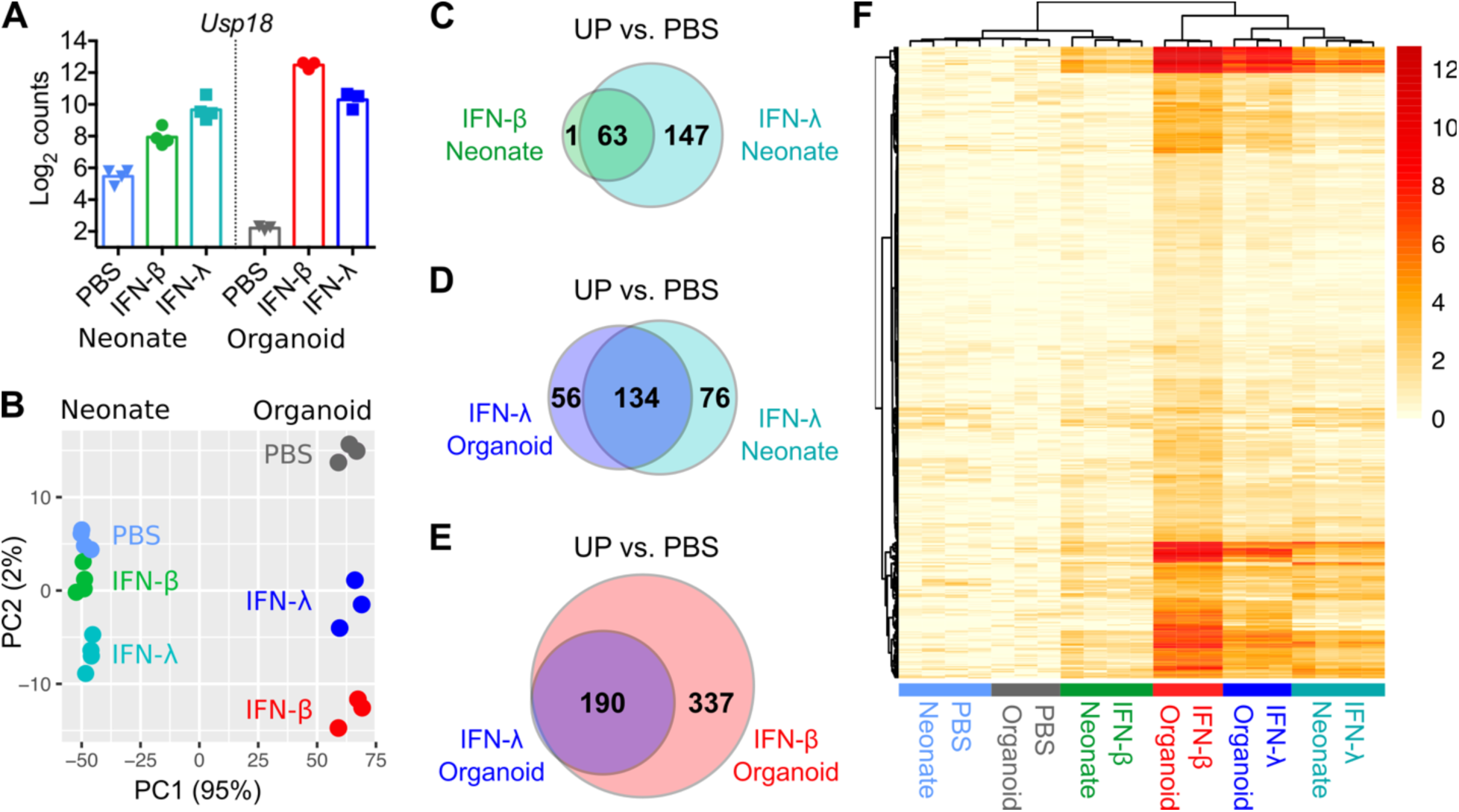
Global ISG expression in response to IFN-β is suppressed *in vivo*. IEC organoids or neonatal mice were treated for four hours with PBS or IFN as in **figures 1-2** and isolated IECs were analyzed by RNAseq. **A**. Normalized read counts for *Usp18*. **B**. PCA analysis of the top 500 differentially expressed genes. **C-E**. Venn diagrams showing the overlap in genes stimulated by the indicated IFN treatment relative to their matched PBS treated controls. **F**. Heatmap comparing log2 fold-change of 527 ISGs among organoid and neonate IFN treatment groups relative to matched PBS controls. Data points represent replicate treatments.

We identified differentially expressed genes among IFN treatment groups relative to their corresponding PBS controls using liberal inclusion criteria (fold-change > 1.5, adjusted p-value < 0.1). Few genes were downregulated by IFN treatments (**Data Set S2, S3**), consistent with the known transcriptional activation downstream of IFN receptors. Identification of ISGs in neonatal IECs revealed 210 IFN-λ-stimulated genes but only 64 IFN-β-stimulated genes (**Fig. 3C, Data Set S2**). Furthermore, all IFN-β-stimulated genes but one (63/64) were present among the 210 IFN-λ-stimulated genes (**Fig. 3C**). Therefore, the global early response of neonatal IECs to IFN-λ is substantially larger than to IFN-β.

IEC organoids had a comparable number of IFN-λ-stimulated genes (190) as neonatal IECs (210), with the majority of genes (134) present in both (**Fig. 3D-E**). However, in striking contrast to neonatal IECs, IEC organoids had a greater number (527) of IFN-β-stimulated genes (**Fig. 3E**). Among IEC organoid treatment groups, there were zero genes unique to IFN-λ, with all 190 IFN-λ-stimulated genes present among the 527 IFN-β-stimulated genes (**Fig. 3E, Data Set S3**). Therefore, the early responses of IEC organoids to IFN-β and IFN-λ are highly overlapping with IFN-λ-stimulated genes comprising a subset of IFN-β-stimulated genes.

To more comprehensively analyze the relationship between IFN responses of neonate and organoid IECs, we normalized the log2 fold-change of all 527 organoid ISGs to matching PBS controls and plotted these changes on a heatmap with hierarchical clustering (**Fig. 3F**). IFN-λ-stimulated neonate and organoid IECs clustered closer to each other than to other treatment groups, supporting the conclusion that IFN-λ responses are similar between neonate and organoid. In contrast, IFN-β-stimulated neonatal IECs clustered closer to PBS controls than to other IFN treatment groups (**Fig. 3F**). These comparisons indicate that the *in vivo* hypo-responsiveness of IECs to IFN-α/β applies globally to all ISGs, whereas responsiveness to IFN-λ is a relatively stable IEC-intrinsic property.

### Apoptosis genes are among IFN-β-specific ISGs

The preceding data from neonatal mice, together with prior studies of adult mice, strongly suggest that hypo-responsiveness of IECs to IFN-α/β *in vivo* is physiologically advantageous. To gain insight into the potential advantages of selective IFN-λ responsiveness, we further analyzed the transcriptomes of IFN-β-responsive IEC organoids. The 337 “IFN-β-specific ISGs” of IEC organoids represented a subset of the overall IFN response that is not present *in vivo* whereas the 190 genes stimulated by IFN-λ and IFN-β represented a “common ISG” module. Notably, the “common ISGs” were stimulated to a significantly greater extent by IFN-β treatment than the “IFN-β-specific ISGs” (**Fig. 4A**), and the 190 common ISGs were stimulated to a significantly greater extent by IFN-β than by IFN-λ treatment (**Fig. 4B**). Therefore, common ISGs consist almost entirely of the most highly responsive genes. To identify differential pathway associations, we compared the 337 IFN-β-specific ISGs with the 190 common ISGs using g:Profiler (33). Comparison of curated pathways from gene ontology (GO), KEGG, and Reactome databases indicated that 1) common ISGs were more significantly associated with antiviral effector pathways, 2) common ISGs and IFN-β-specific ISGs were similarly associated with antigen processing and presentation pathways, and 3) IFN-β-specific ISGs were significantly associated with apoptosis pathways (**Fig. 4C, Data Set S4**). A heatmap of all apoptosis pathway genes (KEGG:04210) confirmed that IFN-β treated IEC organoids clustered separately from other treatment groups and controls (**Fig. 5A**). Specifically, IFN-β treatment of organoids uniquely stimulated 19/130 apoptosis pathway genes including pro-apoptotic genes *Bid, Bcl2l11*, and *Casp8* (**Fig. 5A-B**). Together, these analyses indicate that IFN-β and IFN-λ are similarly capable of eliciting antiviral effectors, but IFN-β uniquely stimulates expression of apoptosis-pathway genes.

**Figure 4.**
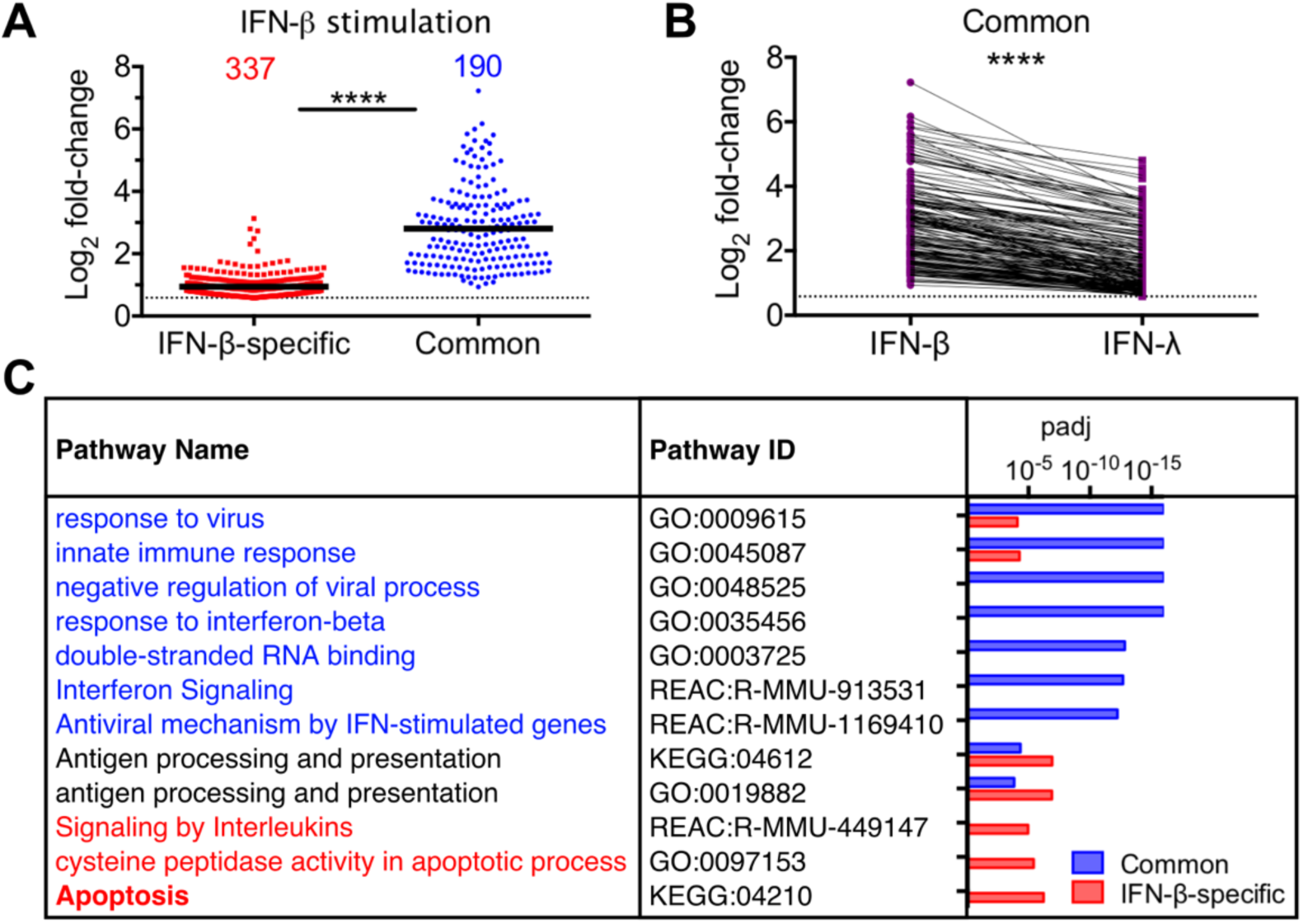
Common ISGs in IEC organoids are highly stimulated and consist of canonical antiviral genes. **A**. Log2 fold-change from RNAseq of genes stimulated by IFN-β but not IFN-λ (337, IFN-β-specific), or genes stimulated by both IFN-β and IFN-λ (190, Common). **B**. Comparison of log2 fold-change for common ISGs following stimulation by IFN-β or IFN-λ. Lines indicated gene identity across treatment groups. **C**. Selected pathways differentiating the indicated ISG categories; “padj” is the p-value adjusted for multiple comparisons. Complete list of pathways in **Data Set S4**. Statistical significance determined by Mann-Whitney test; ****, p < 0.0001.

**Figure 5.**
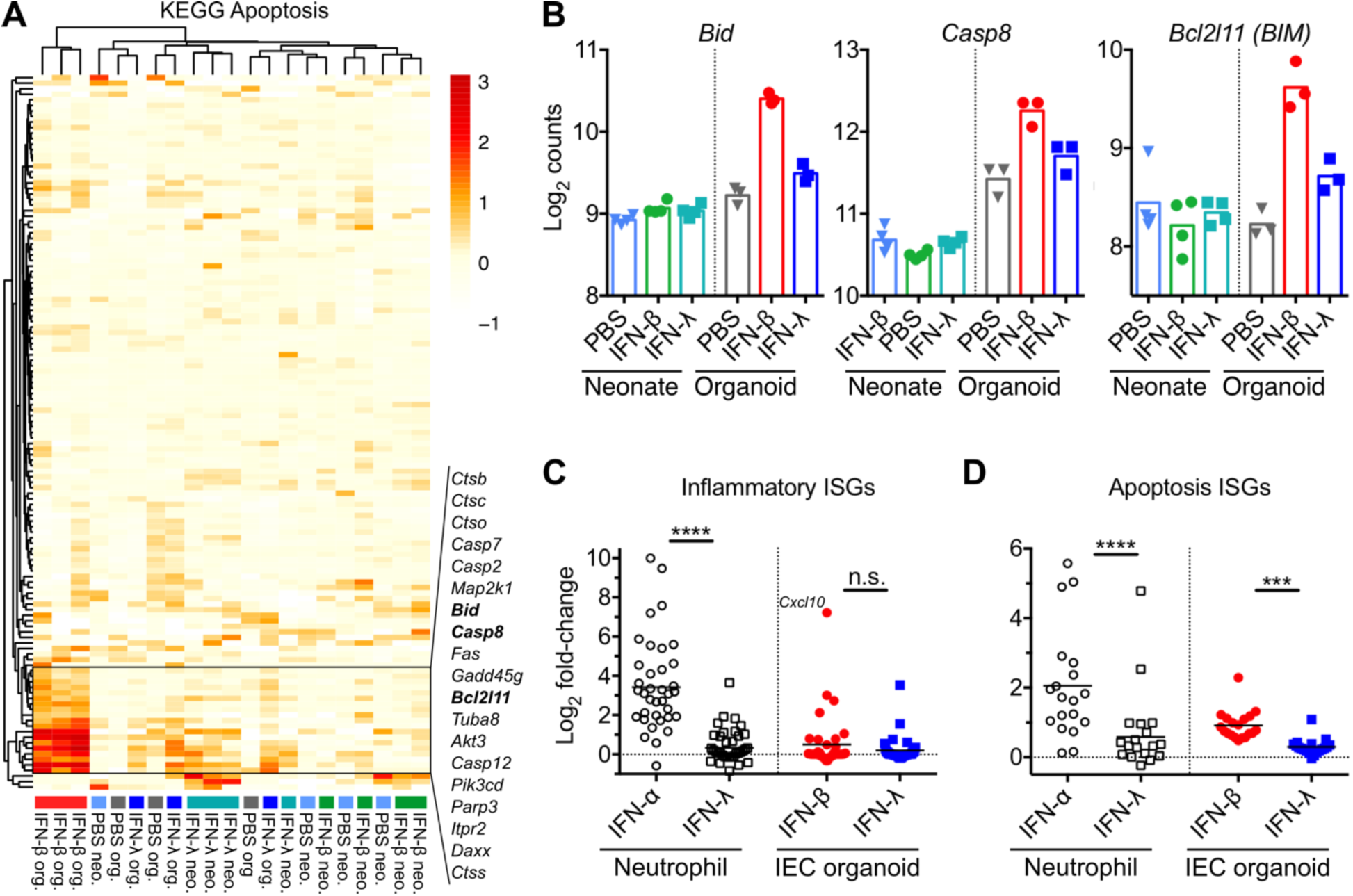
IFN-β-specific ISGs include apoptosis pathway genes. **A.** Heatmap comparing log2 fold change of all KEGG apoptosis pathway genes among organoid and neonate IFN treatment groups relative to their corresponding PBS controls. Names are shown for the cluster of “apoptosis ISGs” stimulated by IFN-β treatment of organoids. **B**. Normalized RNAseq counts for three apoptosis ISGs. **C-D.** Log2 fold change of inflammatory ISGs (**C**) or apoptosis ISGs (**D**) from IFN-treated neutrophils (24) (open circles = IFN-α, open squares = IFN-λ) or IEC organoids (filled circles = IFN-β, filled squares = IFN-λ) relative to their corresponding PBS controls. Statistical significance determined by Kruskal-Wallis test. ***, p < 0.001; ****, p < 0.0001; n.s., p >0.05.

Prior studies in other cell types have indicated that inflammatory cytokines are a gene set that distinguishes IFN-α/β from IFN-λ (22, 24). To determine whether these genes were also differentially regulated in our IEC organoid studies, we performed focused analysis of 37 inflammatory cytokines, including IL-6, IL-1β, and TNFα, shown by Galani et al. to be differentially regulated in neutrophils. Unlike neutrophils, the majority of these inflammatory cytokines were not stimulated in IEC organoids, with the notable exception of the pro-inflammatory chemokine *Cxcl10* (**Fig. 5C, Data Set S5**). This suggests that neutrophils and IECs differ in their capacity for ISG expression. To determine whether this difference extended to the set of IFN-β-specific apoptosis ISGs identified here, we analyzed expression of these genes in the RNAseq data from Galani et al. Similar to our results in IECs, neutrophils upregulated apoptosis pathway genes following treatment with IFN-α but not IFN-λ (**Fig. 5D**). These comparisons suggest that some IFN-α/β-specific ISGs (i.e. inflammatory cytokines) are cell-type specific and others (i.e. pro-apoptotic genes) are similar across cell types.

### Genes stimulated by IFN-β potentiate TNFα-triggered apoptosis

Among apoptosis pathway genes identified in **Figure 5A**, *Bid* and *Casp8* gene products are integral effectors in the extrinsic apoptosis pathway triggered by the inflammatory cytokine TNFα (34). To determine whether IFN-β treatment potentiates TNFα-triggered apoptosis, we pre-treated IEC organoid cultures with IFN-β, IFN-λ, or PBS followed by treatment with TNFα and measured cell viability using the MTT assay (**Fig. 6A**). Treatment with TNFα, IFN-β or IFN-λ alone resulted in minimal loss of IEC viability (<10%) suggesting that our IEC organoids are resistant to the cytotoxic effects of TNFα at baseline. However, IFN-β treatment synergized with TNFα and resulted in an average 30% loss in viability. In contrast, IFN-λ followed by TNFα treatment resulted in significantly less (average 18%) loss of viability (**Fig. 6A**). To confirm that death in these IEC organoid cultures was related to apoptosis, we examined cleaved (active) executioner caspase 3 by immunofluorescence. IFN-β treatment followed by TNFα resulted in a significant increase in the percentage of cleaved caspase 3 in IEC organoids whereas IFN-λ treatment did not (**Fig. 6B-C**). These data indicate that IFN-β stimulation results in greater sensitivity of IECs to TNFα-triggered apoptosis, and suggest that hypo-responsiveness of IECs to IFN-α/β *in vivo* favors epithelial viability.

**Figure 6.**
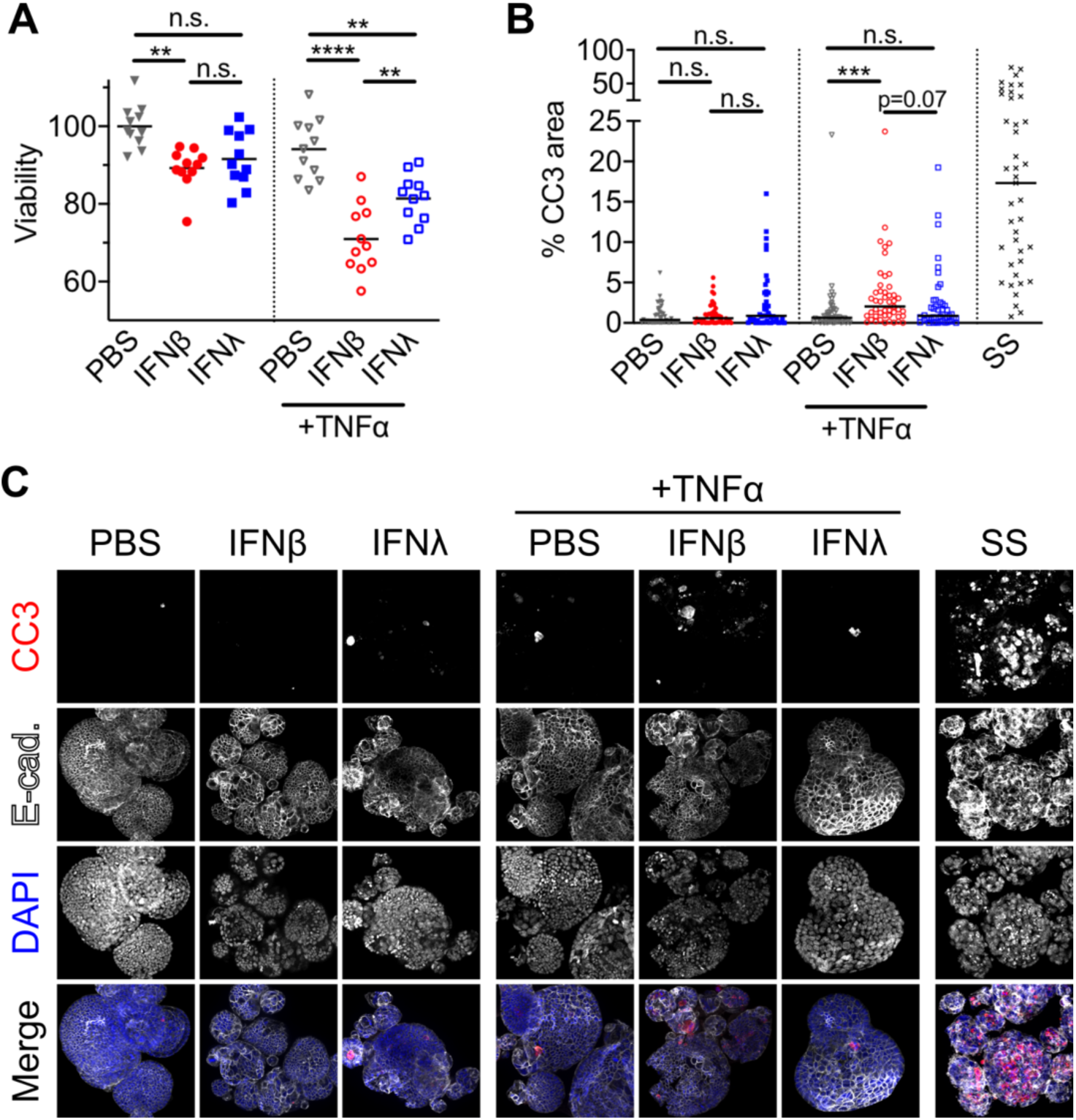
Genes stimulated by IFN-β potentiate TNFα-triggered apoptosis. **A.** MTT viability assay of IEC organoids treated with 10ng/mL IFN-λ3, IFN-β, or PBS for four hours followed by treatment with 100 ng/mL TNFα for 20 hours. **B-C**. Cleaved-caspase 3 (CC3) in IEC organoids was assessed by immunofluorescence following pre-treatment with 10ng/mL IFN-λ3, IFN-β, or PBS for 4-8 hours and subsequent treatment with media or 100ng/mL TNFα for 16-20 hours. The positive apoptosis control, staurosporine (SS), was administered to PBS organoids for 16-20 hours. Data are pooled from three independent experiments with statistical significance determined by one-way ANOVA in **A** and statistical significance determined by Kruskal-Wallis test with Dunn’s multiple comparisons in **B.** Solid line depicts the mean in **A** and the median in **B** with **, p < 0.01; ***, p < 0.001; ****, p < 0.0001; n.s., p >0.05.

### Strength of canonical ISRE determines ISG categories

To globally define distinguishing promoter motifs of common and IFN-β-specific ISG categories, we used the Hypergeometric Optimization of Motif EnRichment (HOMER) software package (35). We searched for motifs enriched in IFN-β-specific ISG promoters relative to a “background” of common ISG promoters or *vice versa*. This comparison resulted in no statistically significant promoter motifs that distinguished IFN-β-specific ISGs from common ISGs. However, a *de novo* motif was significantly enriched in common ISG promoters relative to IFN-β-specific ISG promoters (**Fig. 7A**). This “common ISG motif” was clustered near the transcription start site (TSS) of these genes, consistent with a direct role in initiating transcription (**Fig. 7B**). The *de novo* common ISG motif had a high degree of similarity to previously described ISRE and IRF motifs, suggesting that it reflected stronger canonical promoter motifs among common ISGs. Indeed, previously defined (canonical) ISRE and IRF motifs were present with significantly higher frequency among common ISGs than among IFN-β-specific ISGs (**Fig. 7A, Data Set S6**). Further comparison of common ISGs and IFN-β-specific ISGs to a “background” of other genes within the mouse genome revealed that ISRE and IRF motifs were significantly enriched in both ISG sets, and a STAT1 motif was specifically enriched among common ISGs (**Fig. 7A-B, Data Set S6**). As a control for these comparisons, the GC-rich basal promoter motif of specificity protein 1 (Sp1) was found at a similar frequency near TSS of all gene categories (**Fig. 7A-B**).

**Figure 7.**
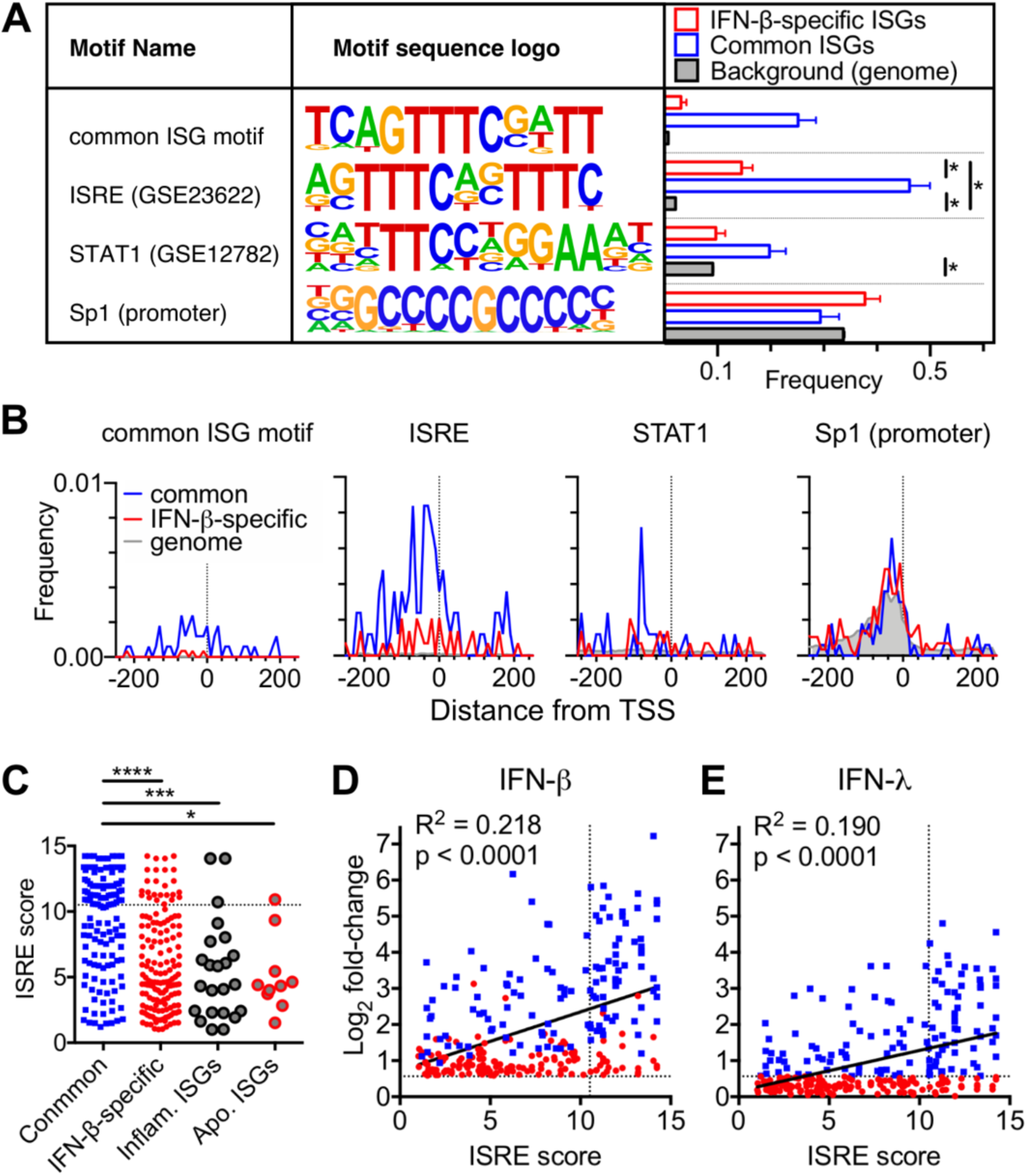
Strength of canonical ISRE differentiates ISG categories. Motifs were identified in genomic sequences 500 bp upstream to 250 bp downstream of annotated transcriptional start sites (TSS) for gene sets. **A**. Motif sequence logos for *de novo* “common ISG motif” and known motifs from the HOMER database. Height of bases are proportional to their preference at that position. Frequency graph depicts the proportion of genes in each category with at least one instance of the indicated motif scoring above threshold, with * indicating q-value < 0.05. **B.** Histograms of motif location relative to TSS. **C.** Comparison of ISRE (GSE23622) motif score among previously defined common, IFN-β specific, apoptosis, and inflammatory ISGs. Dashed line indicates threshold score from analysis in **(A)**; highest scoring motif is shown for each gene with at least one motif score > 1. **D-E**. Correlation of ISRE score with log2 fold-change following treatment with IFN-β (**D**) or IFN-λ (**E**). Vertical dashed line indicates threshold score from analysis in **(A**), horizontal dashed line indicates 1.5-fold differential expression cutoff.

We were interested in determining how well the global analysis of IFN-β-specific ISG promoters reflected properties of apoptosis ISGs and neutrophil inflammatory ISGs. Additionally, we sought to perform more quantitative motif comparisons beyond a simple presence/absence determination. So, we determined the ISRE score for each gene, which is higher for promoter sequences that more closely match the ideal ISRE (**Fig. 7A**). ISRE motif scores were similarly low among apoptosis ISGs, inflammatory ISGs, or IFN-β-specific ISGs, all of which were significantly lower than common ISGs (**Fig. 7C**). These data indicate that promotor characteristics of IFN-β-specific ISGs as a whole are reflected in apoptosis and inflammatory gene subsets.

Prior studies in other cell types have indicated that IFN-λ stimulates less robust STAT1 phosphorylation and ISRE transactivation than IFN-α/β (17, 20). Consistent with these findings, we observed a lower fold-increase of common ISGs by IFN-λ than by IFN-β (**Fig. 4B**). To explore the relationship among ISRE score and fold-increase, we performed correlation analyses of these two variables for all ISGs. We observed a significant positive correlation between IFN-β-stimulated fold-change and ISRE score, confirming the relevance of this promoter motif (**Fig. 7D**). IFN-λ-stimulated fold-change was also significantly correlated with ISRE score, but had a significantly shallower slope (p = 0.0293) (**Fig. 7E**). Together, these data indicated ISGs with low-scoring ISRE motifs were less likely to be stimulated by IFN-λ and correspondingly more likely to be IFN-β-specific.

### Apoptosis ISGs depend on canonical transcription factor STAT1

The preceding analyses suggested that IFN-β-specific ISGs remained functionally dependent on canonical transcription factors in spite of their low-scoring ISRE motifs. To test this experimentally, we generated IEC organoids from *Stat1*^*-/-*^ mice that are unable to generate active STAT1 homodimers, or heterotrimeric ISGF-3. We treated WT and *Stat1*^*-/-*^ organoids with 10ng/mL IFN-β or IFN-λ3 for four hours followed by qPCR to measure induction of common ISGs (*Isg15, Cxcl10*) and IFN-β-specific, apoptosis ISGs (*Casp8, Bid, Bcl2l11*). Common ISGs were induced greater than 1000-fold by IFN-β and greater than 100-fold by IFN-λ in WT IECs. We confirmed that common ISGs were not induced by either IFN type in *Stat1*^*-/-*^ IECs, consistent with an absolute requirement of STAT1 for their stimulation (**Fig. 8A-B**). Pro-apoptotic ISGs were induced two- to three-fold by IFN-β in WT IECs, but were not induced by IFN-λ in WT IECs or by either treatment in *Stat1*^*-/-*^ IECs (**Fig. 8A-B**). These data indicate that the IFN-β-specific ISGs *Casp8, Bid*, and *Bcl2l11* are dependent on canonical ISG transcription factors. Taken together, our findings support the conclusion that IFN-β-specific ISGs are largely distinguished by low-scoring ISRE promoter motifs rather than a unique non-canonical motif. In conclusion, we propose a straightforward model in which a stronger transcriptional response downstream of IFN-β results in an expanded array of ISGs with low-scoring ISRE motifs. These “sub-optimal” ISGs include pro-apoptotic genes that have the potential to disrupt epithelial homeostasis and are therefore physiologically disadvantageous *in vivo*.

**Figure 8.**
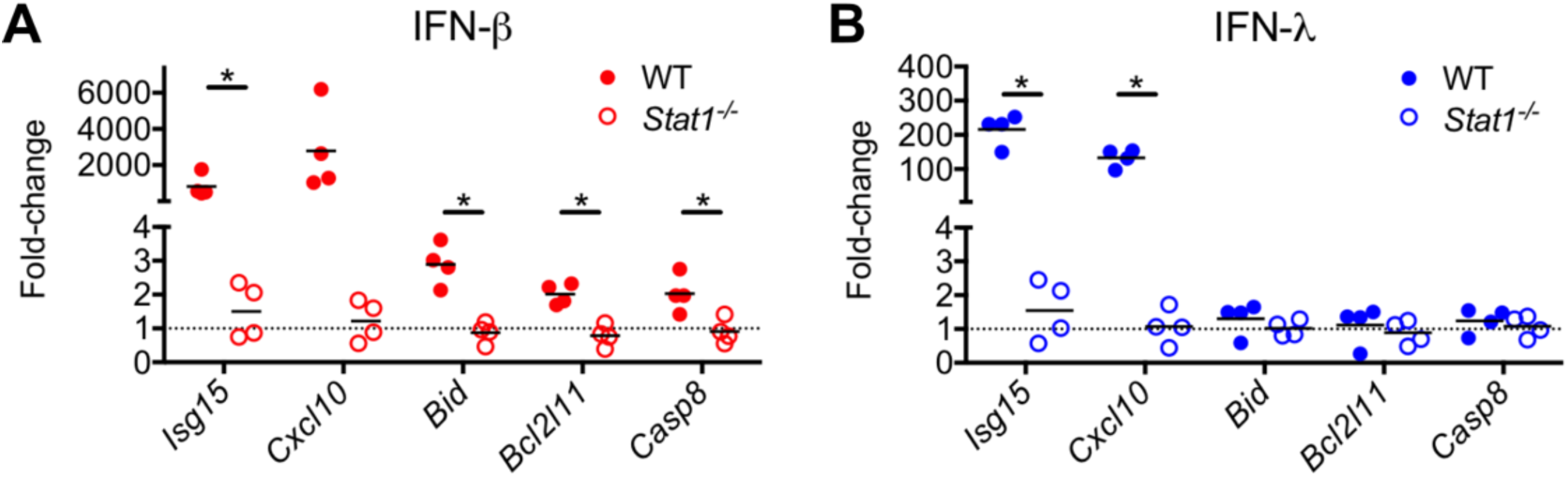
IFN-β-specific ISGs remain dependent on canonical transcription factor STAT1. Quantitative PCR analysis of genes indicated on the x-axis following treatment of WT (filled symbol) or *Stat1*^*-/-*^ (open symbol) IEC organoids with 10ng/mL IFN-β (**A**) or IFN-λ3 (**B**) for four hours; normalized to PBS-treated controls. Data are combined from four experiments, with * indicating p-value < 0.05 by one-way ANOVA.

## DISCUSSION

Here, we find that *in vivo* IECs are hypo-responsive to IFN-α/β beginning in early neonatal life (**Fig. 1**), thereby elevating the importance of IFN-λ in epithelial antiviral immunity. This timing is earlier than anticipated based on the prior study from Lin et al., which found that IFN-α/β stimulated phospho-STAT1 in IECs of neonatal but not adult mice (10). Our transcriptomic comparison of ISG responses indicate that, although a transcriptional response may be marginally stimulated by IFN-β in neonatal IECs, it is significantly diminished relative to the *in vivo* response of neonatal IECs to IFN-λ or the *in vitro* response of IEC organoids to IFN-β (**Fig. 3**). In fact, we find that IEC organoids expanded *in vitro* from intestinal stem cells regain responsiveness to IFN-β, with more robust ISG induction relative to IFN-λ (**Fig. 2**). This IFN response profile of mouse IEC organoids is consistent with recent human IEC organoid studies, which found that IFN-α/β provides more potent antiviral protection from rotavirus (27, 28). We suggest that, similar to mouse IECs, human IECs in their natural context *in vivo* may become hypo-responsive to IFN-α/β.

Our studies of viral infection in mouse IEC organoids here together with prior studies of others in human IEC organoids present a paradox: IFN-λ is the dominant effector in mouse models of gastrointestinal virus infection whereas IFN-α/β elicits superior antiviral defense *in vitro* (27). Our work here shows that IFN-β treatment of IEC organoids, in addition to stimulating significantly greater production of antiviral ISGs than IFN-λ, stimulates an expanded set of “sub-optimal” ISGs (**Fig. 7**). This expanded ISG profile includes pro-apoptotic genes that potentiate TNFα-triggered cytotoxicity (**Fig. 6**). These findings in IECs are reminiscent of recent studies in neutrophils that identified a set of inflammatory cytokine genes, including TNFα, triggered by IFN-β but not IFN-λ (24). We find here that IECs are not capable of readily producing most of these inflammatory ISGs in response to IFN-β, emphasizing the importance of cell-lineage-specific studies (**Fig. 5C**). However, when considered in the context of preceding neutrophil studies, our findings here suggest that a neutrophilic, inflamed intestine is a scenario in which IEC hypo-responsiveness to IFN-α/β would be particularly beneficial. If IECs were not hypo-responsive in this inflammatory scenario, IFN-α/β would synergistically elicit TNFα production by neutrophils and potentiate TNFα-triggered cytotoxicity of IECs. Indeed, such a synergistic response may explain observations of epithelial apoptosis in the IFN-α/β-responsive lung epithelium during influenza infection, where IFNα but not IFN-λ treatment amplifies apoptosis of lung epithelial cells (25). In addition to the less damaging ISG profile of IFN-λ, Broggi et al. showed that IFN-λ can actively suppress inflammatory responses of neutrophils by a post-transcriptional mechanism, providing further homeostatic benefits (26). Thus, hypo-responsiveness of IECs to IFN-α/β *in vivo* may be moderately dis-advantageous for antiviral protection, but reduces the risk of inflammatory response amplification loops that result in epithelial damage.

Previous work has suggested apical trafficking of IFNAR and reduced *Ifnar1/Ifnar2* gene expression as mechanisms for IEC hypo-responsive to IFN-α/β *in vivo* (6, 11). These mechanisms may relate to post-natal changes in intestinal exposure to nutrients and the microbiome, which elicit corresponding changes in IEC metabolism and immunity. Neil et al. recently showed that when the microbiome is depleted from adult mice with antibiotics and the epithelium is exposed to damaging dextran sodium sulfate (DSS), an IFN-α/β response in IECs can promote beneficial recruitment of IL-22-producing leukocytes (36, 37). This role of IFN-α/β in IECs suggests the following possibilities: 1) depletion of the microbiome together with epithelial damage could increase IFN-α/β responsiveness of IECs, or 2) under certain types of epithelial damage the modest responsiveness of IECs to IFN-α/β *in vivo* may be beneficial. Further studies are needed to determine the effect of microbiome and inflammatory triggers on IFN-α/β responsiveness of IECs and to understand the signals that regulate IEC-intrinsic IFNAR responses *in vivo*. Our work here emphasizes the important pleiotropic roles of the IFN response and provides a physiological basis for regulating the ISG expression capacity of IECs to maintain intestinal homeostasis.

## MATERIALS AND METHODS

### Mice

C57BL/6J, BALB/c, and *Stat1*^*-/-*^ (*Stat1*^*tm1Dlv*^) mice were obtained from Jackson Laboratories and bred in specific-pathogen-free barrier facilities at Oregon Health & Science University. Animal protocols were approved by the institutional animal care and use committee at Oregon Health & Science University (protocol # IP00000228) according to standards set forth in the *Animal Welfare Act*. 0.2μg IFN-β (PBL #12405-1) or IFN-λ3 (PBL #12820-1) were administered to seven-day-old neonatal mice via sub-cutaneous injection; an equal volume of diluent (PBS) was administered to littermate control mice.

### Rotavirus infection of mice

Mouse rotavirus, strain EC, was generously provided by Dr. Andrew Gewirtz (Georgia State University). Virus stocks were generated by inoculating four- to six-day-old neonatal BALB/c mice and collecting the entire gastrointestinal tract upon observation of diarrhea four to seven days later. Intestines were subjected to a freeze-and-thaw cycle, suspended in PBS, homogenized in a bead beater using 1.0 mm zirconia/silica beads (BioSpec Products), clarified of debris, and aliquoted for storage at -70°C. 50% shedding dose (SD50) was determined by inoculation of 10-fold serial dilutions in adult C57BL/6J mice. For protection studies, seven-day-old neonatal mice were orally inoculated with 100x SD50 and intestines were isolated 20 hours later for quantitation of viral genomes by qPCR.

### RNAscope

Swiss rolls of intestinal tissue were fixed in 10% neutral-buffered formalin for 18-24hr and paraffin-embedded. Tissue sections (5um) were cut and maintained at room temperature with desiccant until processed. RNA *in situ* hybridization was performed using the RNAscope Multiplex Fluorescent v2 kit (Advanced Cell Diagnostics, ACDBio) per protocol guidelines. Staining with anti-sense probes for detection of *Usp18* (ACDBio #524651) was performed using ACDBio protocols and reagents. Slides were stained with DAPI and mounted with ProLong Gold antifade reagent (ThermoFisher), and imaged using a Zeiss ApoTome2 on an Axio Imager, with a Zeiss AxioCam 506 (Zeiss).

### Organoid culture

Primary organoid culture media was Advanced DMEM/F12 (ThermoFisher #12634010) supplemented with 20% fetal bovine serum, 1X penicillin/streptomycin/L-glutamine, and 10mM HEPES. Isolation and culture of primary mouse IEC organoids was performed essentially as described (38). Briefly, intestinal crypts were isolated by mechanical disruption, digestion with 2mg/mL collagenase type I, and centrifugation. Isolated crypts were resuspended in 15uL Matrigel (Corning #354234) per well and plated in 24-well plates. Organoids were grown and maintained in 50% primary organoid culture media mixed with 50% conditioned media (CM) from L-WRN cells (ATCC cat#CRL-3276), which contained Wnt3a, R-spondin3, and Noggin. ROCK-inhibitor (Selleck Chemicals #S1049) and TGFbeta-inhibitor (Selleck Chemicals #S1067) were added to culture media to promote survival of dissociated cells. Media was replaced every two days. Every three days, or when organoids become dense, cells were disrupted with Trypsin/EDTA and replated at ∼30,000 cells/well. IFN-β (PBL #12405-1), pegylated-IFN-λ2 (Bristol Meyers-Squibb), IFN-λ3 (PBL #12820-1), and TNFα (Peprotech #315-01A) were added to organoid cultures as indicated in the figure legends.

IEC organoid viability assays were adapted from Grabinger et al (39). Organoids were seeded in 96 well plates at 500-1000 cells per 5 uL matrigel per well. Following cytokine treatment as indicated in figure legend, MTT (Sigma #MM5655) was added to cell culture media at 0.5 mg/mL and incubated at 37 °C for 1 hour. Media was replaced with 100% DMSO and absorbance was measured at 570 nm on a BioTek plate reader. Background-subtracted OD values were normalized to untreated organoid wells (100% viability) for each independent experiment.

### Rotavirus infection of organoids

Primary mouse IEC organoids were dissociated to the single-cell level with Trypsin/EDTA and were seeded at ∼25,000 cells/well. Organoids were maintained in 50% CM for two days followed by one day of culture in 5% CM – ROCK inhibitor and TGFbeta inhibitor were supplemented to maintain the concentrations present in 50% CM. Cells were treated for 8-hours with 10ng/mL IFN-β (PBL #12405-1), 10ng/mL IFN-λ3 (PBL #12820-1), or PBS control. Organoids were inoculated with 500x SD50 murine rotavirus EC in 5% CM by overlaying inoculant, rocking at room-temperature for 30 minutes, and washing with PBS three times. Infected cells were incubated in 5% CM until the indicated timepoints. Cells were lysed in ZR Viral RNA Buffer (Zymoresearch) and viral genomes were detected by quantitative RT-PCR.

### Organoid immunofluorescence

Following stimulation, organoids were removed from Matrigel by rocking at 4°C in cell recovery solution (Corning). Cells were stained by adapting previous protocols (40). In short, cells were fixed in 3.7% paraformaldehyde, permeabilized in ice-cold 100% methanol, and blocked in 5% normal goat serum, 5% bovine serum albumin, and 0.5% saponin in PBS. Cells were stained with mouse anti-Ecadherin (Becton Dickinson, #610182), rabbit anti-cleaved caspase 3 (Cell Signaling Technology, #9661S), secondary goat anti-mouse Alexa Fluor 555, and goat anti-rabbit Alexa Fluor 647 (ThermoFisher) in 1% normal goat serum, 1% bovine serum albumin, and 0.5% saponin in PBS. IEC organoids were counterstained with DAPI, mounted with ProLong Gold antifade reagent (ThermoFisher), and imaged using a Zeiss ApoTome2 on an Axio Imager, with a Zeiss AxioCam 506 (Zeiss). The CC3-positive area was measured and normalized to the total area of organoid surface using ImageJ.

### Quantitative RT-PCR

RNA was isolated using RiboZol (Amresco) or the ZR Viral RNA Kit (Zymoresearch). DNA contamination was removed using the DNAfree kit (Life Technologies). cDNA was generated with the ImPromII reverse transcriptase (Promega). Quantitative PCR was performed using PerfeCTa qPCR FastMix II (QuantaBio) and the following primers and probes: *Bid* – Integrated DNA technologies (IDT) assay #Mm.PT.58.8829163; *Bcl2l11* – IDT assay #Mm.PT.58.12605058; *Casp8* – IDT assay #Mm.PT.58.41467226; *Isg15* – IDT assay #Mm.PT.58.41476392.g; *Usp18* – IDT assay #Mm.PT.58.28965870; *Cxcl10* – IDT assay #Mm.PT.58.28790444; *Rps29* – IDT assay Mm.PT.58.21577577; Mouse Rotavirus – Primer1: GTTCGTTGTGCCTCATTCG, Primer2: TCGGAACGTACTTCTGGAC, Probe: AGGAATGCTTCAGCGCTG. Absolute copy number was determined by comparing Ct values to a standard curve generated using DNA of known copy number encoding the target sequence. Samples are graphed as absolute copy number of the indicated target divided by absolute copy number of the housekeeping gene (*Rps29*). Samples with fewer than 1,000 copies of housekeeping gene were excluded.

### Cell isolation and flow cytometry

Epithelial fractions were prepared by non-enzymatic dissociation as previously described (41). Briefly, mouse intestines were isolated, opened longitudinally, and incubated in stripping buffer (10% bovine calf serum, 15 mM HEPES, 5 mM EDTA, 5 mM dithiothreitol [DTT] in PBS) with shaking for 20 min at 37°C. The dissociated cells were collected and stained for fluorescence-activated cell-sorting (FACS). Cells were stained with the Zombie Aqua viability dye (Biolegend), Fc receptor blocking antibody (CD16/CD32; Biolegend), anti-EpCAM (clone G8.8; Biolegend), and anti-CD45 (clone 30-F11; Biolegend). Cells in the live gate were sorted as EpCAM-positive/CD45-negative IECs or EpCAM-negative/CD45-positive hematopoietic cells.

### RNA sequencing

Quality of RNA samples were assessed using a TapeStation (Agilent) and mRNA sequencing libraries were prepared using the TruSeq RNA Library Prep Kit (Illumina). Barcoded triplicate samples from IEC organoids (9 total) and quadruplicate samples from neonates (12 total) were separately prepared and pooled. Single-read sequencing was performed using an Illumina HiSeq 2500 through the Massively Parallel Sequencing Shared Resource at OHSU.

### Gene expression analysis

Adaptor-trimmed reads were mapped to the mouse genome (GRCm38) using the STAR aligner (42), and mapping quality was evaluated using RSeQC (43), and MultiQC (44). All samples had between 15 and 30 million uniquely mapped reads with similar distribution across genomic features and uniform gene-body coverage. Read counts per gene were determined using the featureCounts program (45), and differential expression analysis was performed using DEseq2, as described (46). PCA analysis was performed on DEseq2 regularized logarithm (rlog) transformed data. Heatmaps were generated using log2-transformed data normalized to the mean of matched PBS control samples; heatmap clustering is based on Euclidean distance.

### Statistical Analyses

Data were analyzed with Prism software (GraphPad Prism Software), with specified tests as noted in the figure legends.

### Data availability

RNA sequencing data obtained in this study have been deposited in the NCBI Gene Expression Omnibus under GEO Series accession number GSE142166.

## Supporting information

Data Set S1

Data Set S2

Data Set S3

Data Set S4

Data Set S5

Data Set S6

Data Set S7

## Acknowledgments

The authors would like to thank the following OHSU core facilities for technical support: Integrated Genomics Laboratory, Advanced Light Microscopy Core, Flow Cytometry Core, and Histopathology Core. T.J.N. was supported NIH grant R01-AI130055 and by a faculty development award from the Sunlin & Priscilla Chou Foundation. J.A.V. was supported by NIH grants T32-GM071338 and T32-AI007472. D.A.C was supported by NIH grant T32-AI007472. The funders had no role in study design, data collection and interpretation, or the decision to submit the work for publication.

## Author Contributions

Conceptualization, T.J.N.; Methodology, J.A.V., D.A.C., L.L., and T.J.N.; Investigation, J.A.V., D.A.C., L.L., and T.J.N.; Writing – Original Draft, T.J.N.; Writing – Review & Editing, J.A.V. and D.A.C.; Visualization, T.J.N.; Supervision, T.J.N.; Project Administration, L.L. and T.J.N.; Funding Acquisition, T.J.N.

## Disclosure

We declare no competing interests.

## SUPPLEMENTAL FILES

**Data Set S1.** Differentially expressed genes in IECs of neonates and organoids. Tab 1 is the DESeq2 output list of genes differentially expressed in neonate IECs relative to organoid IECs cut-off at padj <0.05. Tab 2 lists genes from tab 1 with a log2 fold-change (l2fc) < -1 (DOWN) or l2fc > 1 (UP). Tab 3 lists pathways significantly associated with DEGs UP in neonate relative to organoid. Tab 4 lists pathways significantly associated with DEGs DOWN in neonate relative to organoid.

**Data Set S2.** Differentially expressed genes in IECs of neonatal mice treated with IFN. Tab 1 is the DESeq2 output list of genes with differential expression following IFN-β treatment (nIFNB_vs_nPBS) cut-off at padj < 0.1. Tab 2 is the DESeq2 output list of genes with differential expression following IFN-λ (nIFNL_vs_nPBS) cut-off at padj < 0.1. Tab 3 lists genes from prior tabs with greater than 1.5-fold increase (l2fc > 0.585) as depicted in **figure 3C**.

**Data Set S3.** Differentially expressed genes in IEC organoids treated with IFN. Tab 1 is the DESeq2 output list of genes with differential expression following IFN-β (IFNBvsPBS) cut-off at adjusted p-value (padj) < 0.1. Tab 2 is the DESeq2 output list of genes with differential expression following IFN-λ (IFNLvsPBS) cut-off at padj < 0.1. Tab 3 lists common ISGs and IFN-β-specific ISGs as defined in **figure 3E**.

**Data Set S4.** Pathways of differentially expressed genes in IEC organoids treated with IFN. A complete list of pathways with at least one gene present from common ISGs (tab 1, Common_Pathways) and IFN-β-specific ISGs (tab 2, IFNB_Pathways). Highlighted rows indicate the curated set of pathways shown in **figure 4A**.

**Data Set S5.** Re-analysis of gene sets in neutrophils treated with IFN for four hours from Galani et al. Tab 1 is the DESeq2 output list of genes with differential expression following IFN-α (neutIFNA_vs_neutUNTX) cut-off at padj < 0.1. Tab 2 is the DESeq2 output list of genes with differential expression following IFN-λ (neutIFNL_vs_neutUNTX) cut-off at padj < 0.1. Tabs 3 and 4 list log2 fold-changes in neutrophils and IEC organoids for the set of previously defined inflammatory genes (tab 3) and the set of apoptosis genes defined in **figure 5A** (tab 4).

**Data Set S6.** Enrichment of known motifs within the Homer database among common ISGs relative to a background of IFN-β-specific ISGs (tab 1, common_vs_infb), IFN-β-specific ISGs relative to a background of common ISGs (tab2, ifnb_vs_common), common ISGs relative to a background of the mouse genome (tab 3, common_vs_genome), and IFN-β-specific ISGs relative to a background of the mouse genome (tab 4, ifnb_vs_genome). Bold rows indicate associations with a q-value of < 0.05. Highlighted rows indicate motifs depicted in **figure 8**.

**Data Set S7.** ISRE motif scores for the indicated genes; related to **figure 7C-E**. Only highest scoring ISRE motifs for each gene are shown. Only genes with at least one motif score of > 1 are shown. Top ISRE score for all 527 organoid ISGs were compared to log2 fold-change from RNAseq (tab 1, ISREscore_vs_log2fc_all_ISGs). Top ISRE scores were also calculated for inflammatory ISGs (tab 2, ISREscore_Inflammatory_ISGs) and apoptosis ISGs (tab 3, ISREscore_apoptosis_ISGs) as depicted in **figure 7C**.

## Notes

https://www.ncbi.nlm.nih.gov/geo/query/acc.cgi?acc=GSE142166

